# Computational Study of Ion Permeation Through Claudin-4 Paracellular Channels

**DOI:** 10.1101/2022.03.17.484760

**Authors:** Alessandro Berselli, Giulio Alberini, Fabio Benfenati, Luca Maragliano

## Abstract

Claudins (Cldns) form a large family of protein homologs that are essential for the assembly of paracellular tight junctions (TJs), where they form channels or barriers with tissue-specific selectivity for permeants. In contrast to several family members whose physiological role has been identified, the function of claudin 4 (Cldn4) remains elusive, despite experimental evidence suggesting that it can form anion-selective TJ channels in the renal epithelium. Computational approaches have recently been employed to elucidate the molecular basis of Cldns’ function, and hence could help in clarifying Cldn4 role. In this work, we use structural modeling and all-atom molecular dynamics simulations to transfer two previously introduced structural models of Cldn-based paracellular complexes to Cldn4, in order to reproduce a paracellular anion channel. Free energy (FE) calculations for ionic transport through the pores allow us to establish the thermodynamic properties driving the ion-selectivity of the structures. While one model shows a cavity permeable to chloride and repulsive to cations, the other forms barrier to the passage of all the major physiological ions. Furthermore, our results confirm the charge selectivity role of the residue Lys65 in the first extracellular loop of the protein, rationalizing Cldn4 control of paracellular permeability.

## INTRODUCTION

Claudins (Cldns) form a family of 27 homologous proteins with a pivotal role in endothelial and epithelial tight junction (TJ)^1–6^ structure and function. Available structural information show that Cldns fold in a four-helices bundle (named from TM1 to TM4) that embeds in the cellular membrane and anchors two extracellular loops (ECL1-2) and a loop in the cytoplasmic region, where the two terminal domains are also contained. Cldns assemble in TJs via: (i) the formation of strands of protomers lining on the same cell membrane, held together by *cis* interactions, and (ii) intercellular aggregates involving *trans* interactions between the ECLs belonging to the opposite protomers, which seal the thin layer separating two neighboring cells, named the paracellular space. The ECLs arrange in a *β*-sheet layer and, typically, ECL1 affects the TJ permeability, while ECL2 determines the *trans* interactions between Cldns belonging to two adjacent cells^7^. These proteins regulate the paracellular transport of ions and molecules via highly selective mechanisms^8,9^, and for this reason their dysfunction is directly associated with clinical disorders^10,11^. While the physiological role of few Cldns has been extensively characterized^6,7^, others are still to be completely understood. A remarkable example is the claudin 4 member (Cldn4), highly expressed in the kidney but also present in other tissues^12–19^. Recent investigations suggest that Cldn4 may play a relevant role in the dynamics of tumor growth, and its attitude to bind the *Clostridium perfringens* enterotoxin (CPE), as a receptor, can open new approaches for drug delivery strategies^20–22^. Several studies suggest that Cldn4 is responsible for anion reabsorption in the collecting ducts, forming highly selective channels for chloride^23–27^, most likely due to a positively charged residue, Lys65, belonging to the ECL1 domain^23,28,29^. However, the ability of Cldn4 to form TJ strands has been found to depend on the cellular system in which it is expressed^26,30–32^. While biochemical data reported by Hou et al.^23^ suggest that heterotypic interactions between Cldn4 and Cldn8 are required for a functional localization of the TJ in the kidney epithelial cells, homotypic Cldn4 interactions have also been observed^33^.

In the last years, structural modeling and molecular dynamics (MD) simulations, have become instrumental to grasp the fine details of biophysical processes, such as protein-protein aggregation, protein conformational transitions and selectivity of ion channels^34^. More recently, MD simulations allowed the atomic description of the structural and functional features of various Cldn-based paracellular aggregates, including claudin 15 (Cldn15), claudin 5 (Cldn5), and claudin 2 (Cldn2)^35–40^. A remarkable step toward a detailed description of TJ proteins was the introduction of the first model for Cldn15-based channels in Ref. 41, often referred to as “Suzuki model”. The proposed arrangement is formed by multiple copies of the crystal structure of the isolated Cldn15 monomer (PDB ID: 4P79)^42^ and it is consistent with cys-scanning mutagenesis experiments and freeze-fracture microscopy imaging. In this multimeric assembly, linear strands of neighboring Cldns are held together by *cis* interactions. Strands between two opposite cells seal the paracellular space and the ECLs of opposite *cis* dimers form *β*-barrel super-secondary structures that result in pores of radius smaller than 5 Å. Subsequent computational works contributed to refine and validate the original Cldn15-based model^37,38,40,43^, also extending the pore configuration to channels of other members of the Cldn family and assessing its consistency with the tissue-specific physiological properties^36^. A second model of a Cldn-based pore was suggested in Refs. 35,39. Accordingly, two protomers belonging to the same cell assemble in a dimer via interactions between the TM2 and the TM3 helices^44^. Focusing on Cldn5, the authors emphasized that the structural stability of the dimer is due to the formation of a *leucine zipper* involving residues Leu83, Leu90, Leu124 and Leu131, supported by two homophilic *π-* π-interactions between the aromatic residues Phe127 and Trp138 on the opposing TM domains. This specific arrangement for Cldn5 protomers matches the experimental results by Rossa et al.^45^. As described in the computational investigation of Refs. 35,39, two copies of these dimers, belonging to two opposing cells, can associate across the paracellular space to form another pore configuration. Coarse-grained (CG) self-assembly simulations suggested that this second model could be suitable for the assembly of Cldn4-based TJs^35^ as well, but the application to other homologs is highly questionable due to experimental evidences^46^.

In this work, we present a computational investigation of Cldn4 in the two pore arrangements, in order to assess their transferability to this member of the Cldn family. Consistently with the notation of Refs. 35,39,44, we refer to the two models described above as Pore I and Pore II, respectively. The two tetramers are simulated in explicit double membrane bilayer and water environments, and free energy (FE) calculations are performed for single ion or water molecule permeation through the paracellular space. Results show that the two models are permeable to water with no relevant FE barriers for both the pores. Pore I is attractive for chloride, since its energetic profile displays a minimum of −2.5 kcal/mol in the inner portion of the cavity, where the Lys65 residues^23^ belonging to the four protomers form a cage that functions as a selectivity filter. Indeed, the passage of cations is prevented by barriers of ~4 kcal/mol and ~8 kcal/mol for monovalent and divalent cations, respectively, with peaks framed by the same basic residues. In contrast, due to the drastically different distribution of charged pore-lining residues, Pore II is impermeable to all the tested ions, showing double FE barriers of ~3 kcal/mol and ~6 kcal/mol for monovalent and divalent cations, respectively, and a single barrier of ~5 kcal/mol for the anion.

These results reveal that the specific pore arrangement can strongly affect the selectivity properties of Cldn-based TJ pores. In particular, the thermodynamic features of ionic permeation in the different Cldn4 pore models suggest that only the Pore I structure is consistent with the anionic selectivity of Cldn4 paracellular pores.

## METHODS

### Pore I assembly

Pore I was assembled with four Cldn4 monomers, following the protocol illustrated in Ref. 37 and matching the structure published by Suzuki et al.^41^. The high-resolution Cldn4 structure (PDB ID: 5B2G)^47^ is unlikely to be representative of the protein native conformation, because of its binding to the C-terminal fragment of the *Clostridium perfringens* enterotoxin. For this reason, the Cldn4 monomers were homology modelled starting from the Cldn15 structure, available in the isolated form (PDB ID: 4P79)^42^. The SWISS-MODEL platform^48,49^ was used to generate the starting Cldn4 configuration. Subsequently, the model was refined with ModRefiner^50^. The resulting protomer was replicated in four units and superimposed on the template of the Suzuki model^41^ using the UCSF Chimera^51^ Matchmaker tool. Afterwards, a further refinement of the system was performed with GalaxyRefineComplex^52,53^. The *cis*-interactions occurring between two protomers in the same cell are shown in **Fig. 1A**, and the tetrameric structure sealing the paracellular space is introduced in **Fig. 2A,B**.

**Figure 1.**
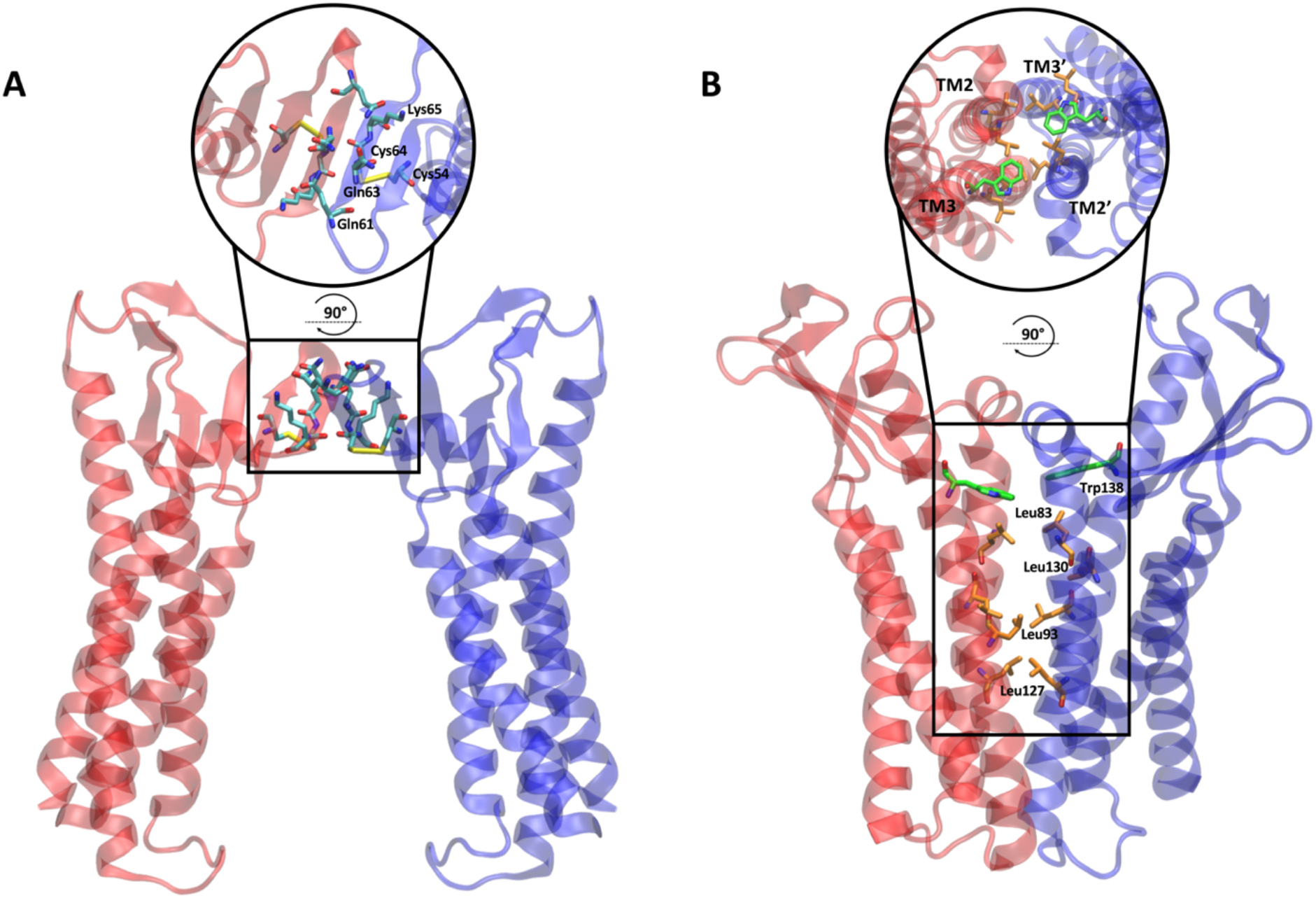
Dimeric cis-interfaces of the two pore models. **A** In the Pore I model, two protomers of the same cell interact at the level of their ECLs, resulting in a hydrophilic interface. The apical zoomed view of the interface is shown in the circle. **B** In the Pore II model, two protomers of the same cell interact at the level of the TM domain forming a hydrophobic interface made by a leucine zipper involving the Leu83, Leu130, Leu93 and Leu127 residues and supported by the π - π interactions between the aromatic Trp138 sidechains. The apical view of the interface is shown in the circle.

**Figure 2.**
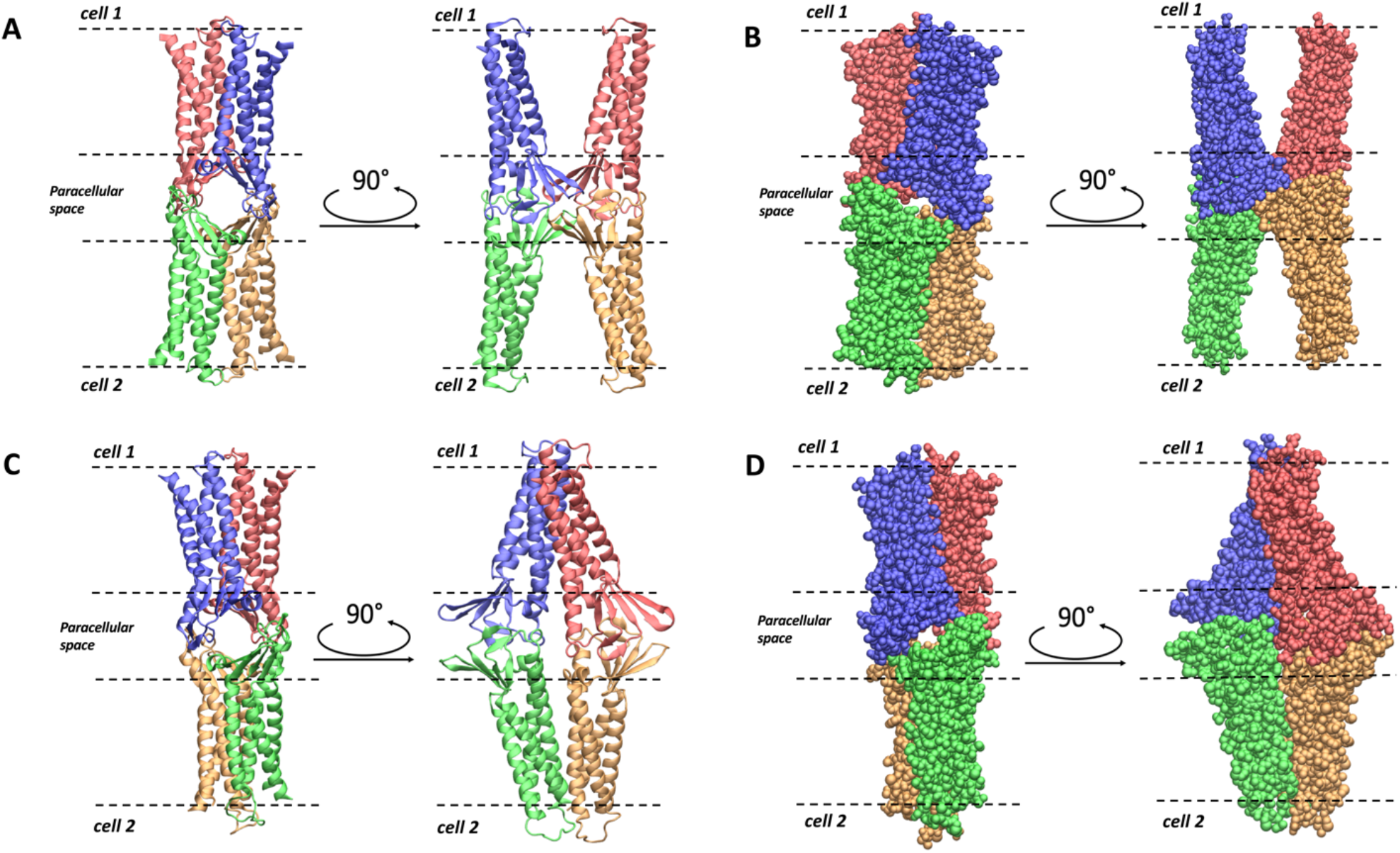
Representations of the Pore I (A,B) and Pore II (C,D) models. **A** *Apical/basolateral and lateral view of the ribbon representation of the Pore I structure*. **B** *Apical/basolateral and lateral view of the Van der Waals representation of Pore I*. **C** *Apical/basolateral and lateral view of the ribbon representation of the Pore II structure*. **D** *Apical/basolateral and lateral view of the Van der Waals representation of Pore II. The four Cldn4 protomers are distinguished by their coloring. In all the panels, the dashed lines identify the boundaries of the membranes of two opposing cells separated by the paracellular space*.

### Pore II assembly

Because of the lack of a reference model based on experimental data of the pore, the strategy applied to reproduce the tetrameric architecture of the Pore II configuration was different from the one adopted for Pore I. First, we generated and equilibrated the *cis-*dimer reproducing the leucine zipper (**Fig. 1B**). A Cldn4 dimer with the specific protein-protein interface was obtained using MEMDOCK^54^. Following an additional refinement with the Rosetta DOCKING2 tool^55–57^, the structure was equilibrated in a homogeneous 1-palmitoyl-2-oleoyl-sn-glycero-3-phosphocholine (POPC) membrane solvated with explicit three-point (TIP3P)^58^ water molecules and neutralized with a physiological KCl concentration and ~250 ns of all-atom MD simulation were performed with the NAMD 3.0 program^59^. The CHARMM36m force field^60^ was used, including the associated ionic parameters with the NBFIX corrections^61–63^. Then, ClusPro^64–68^ was used to build the tetramer, starting from two copies of the equilibrated dimer. Positional restraints were included to keep the TM domains of the opposing dimers far from each other and to ensure the *trans*-interactions of the ECLs domains. Afterwards, the structure was relaxed using the GalaxyRefineComplex server^52,53^ (**Fig. 2C,D**).

### Double Bilayer set-up

Each tetrameric system was embedded in a double POPC bilayer, solvated with explicit three-point (TIP3P)^58^ water molecules and neutralized with counterions, as described below. The pore axis was oriented along the VMD^69^ y-axis. The CHARMM pdb file of the protein complex was generated with the CHARMM-GUI PDB manipulator tool^70,71^. A single hexagonal POPC bilayer, inscribed in a 128.0 × 128.0 Å square, was provided by the *membrane builder* tool of the same platform^71,72^. Two equilibrated copies of the membrane, based on NAMD simulations with the CHARMM36 force field^73^, were used to embed the pore transmembrane domains. After the removal of steric clashes between proteins and lipids, the resulting hexagonal box was filled with explicit TIP3P^58^ water. When needed, counterions were added in both the cytosolic and the paracellular layers. The 4 disulfide bonds between the Cys54 and Cys64 residues were preserved.

#### Equilibration and unbiased MD simulation

After an initial short energy minimization, 30 ns of equilibration were performed by progressively releasing positional restraints on the heavy atoms. The system was simulated in the NPT ensemble at T = 310 K and P = 1 bar, maintained by a Langevin thermostat, with a damping coefficient of 1 ps, and a Nosé-Hoover Langevin piston, adopting an oscillation period of the piston of 50 fs and a damping time scale at 25 fs, respectively^74,75^. The NAMD 3.0 program^59^ in combination with CHARMM36m force field^60^ was used. The hexagonal box was inscribed in a square of about 128.0 × 128.0 Å and of height around 160.0 Å. Periodic boundary conditions were used to replicate the system and remove surface effects. Long-range electrostatic interactions were computed using the Particle Mesh Ewald (PME) algorithm^76^, with an order 6 spline interpolation and a maximum space between grid points of 1.0 Å. Short range electrostatic and van der Waals interactions were calculated with a 12 Å cutoff and using a smoothing decay starting to take effect at 10 Å. A 16 Å pairlistdist was chosen for the neighbor list, and a 2 fs time step was employed. Chemical bonds involving hydrogen atoms and protein heavy atoms were constrained with SHAKE^77^, while those of the water molecules were kept fixed with SETTLE^78^.

The transmembrane domain of each protomer was held fixed by harmonically restraining the *Cα* atoms of residues 11, 14, 25, 28, 78, 81, 99, 102, 116, 119, 143, 146, 166, 169, 183, 186 to their starting positions. Additional harmonic restraints on the *Cα* atoms of residues 30, 35, 40, 45, 50, 55, 60, 65, 70, 150, 153, 156, 159, belonging to the ECLs, were added to Pore II during the production phase. The use of restraints on these atoms mimics the constriction exerted on the structure by the neighboring protomers in the TJ strand, that are not included in the single-pore MD simulations. For each of the two systems, 250 ns of standard MD simulations were performed as additional equilibration, maintaining only the restraints of the *Cα* atoms. The final configurations were adopted for the free energy (FE) calculations.

##### Pore size analysis

The size of the paracellular channel was monitored along the trajectory with the HOLE program^79,80^. This algorithm maps the radius of a channel along a given axis (here, the paracellular channel was oriented along the VMD y-axis) by fitting a spherical probe with the Amber Van der Waals radii^81^ of the pore-lining atoms. A threshold of 10 Å was chosen to define the boundaries of the channel.

#### Free energy calculations

Each FE profile was calculated with the Umbrella Sampling (US) method^82^, where a restraining potential term is added to the MD potential to confine a collective variable (CV) in selected regions, named *windows*, allowing proper sampling also of the high-energy regions. Here, the CV is represented by the coordinate of the tagged permeating ion/water molecule along the pore axis, which was oriented along the VMD Cartesian y-axis. The restraining potential *V_i_*(*y*) in each window *i* is:

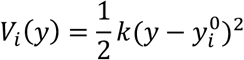

where 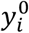 indicates the value in Å at which the CV is restrained in the window (called *center*) and *k* is a constant that is appropriately chosen in order to ensure a sufficient overlap of the CV distributions of adjacent windows (in this work, we used *k* = 2.0 *kcal*/(*mol* Å^2^) for all the simulations). In each window, the displacement of the ion orthogonal to the pore axis is confined within a disk of radius *r*_0_ + *δ*, where *r*_0_ is the pore radius as determined by the HOLE program^79,80^ and *δ* = 2 Å. The equilibrated conformation of the system was used as the starting structure of all the US windows, and the ion/water molecule was manually positioned at each center 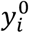. The Pore I channel axis was split into 60 windows spaced 1 Å from each other. After an initial minimization, 16 ns-long trajectories were produced using the same set-up and parameters of the unbiased MD simulation. The first 1 ns was excluded from the analysis for all the windows. Because of its elongated shape, sampling the Pore II model required 75 1-Å-spaced windows. The minimization, equilibration and production procedures followed the same protocol of the Pore I MD run and up to 20 ns per window were simulated in order to achieve proper convergence of the FE profiles. The FE landscapes were obtained adopting the *weighted histogram analysis method* (WHAM)^83–85^ using the code available at http://membrane.urmc.rochester.edu/content/wham^86^, which calculates the statistical error associated to the FE estimation using the bootstrap method. The CV values were written using a 10 ps frequency.

#### Electrostatic potential surface

The electrostatic potential surface was computed with the Adaptive Poisson-Boltzmann Solver (APBS) code^87^, using the default parameters set by the developers. Relative dielectric constants of 2 and 78.54 used for the protein and the solvent, respectively, and the calculations were performed at a temperature of 298.15 K. The surface is shown with a red-white-blue color map ranging from −5 to +5 kT/e.

## RESULTS

By comparing the two Cldn4-based pore models, the position of the amino acids along the channel axis changes completely as a result of the opposite relative orientations of the monomers (**Fig. 3**). In Pore I, we observe two pairs of ECL1 glutamine residues (Gln61 and Gln63) facing each other in the middle of the channel, with their sidechains pointing toward the lumen (**Fig. 3A,B**). Moving away from the center along the axis, two pairs of Lys65 are found next to the glutamines, followed by other acidic (Asp48, Asp68 and Asp146) and basic (Arg31 and Arg158) residues. In Pore II, the positions of all these amino acids are inverted, with the glutamines at the mouths and the charged residues towards the central region (**Fig. 3C,D**). In our MD simulation, the Pore I model revealed a remarkable stability of the paracellular *β*-barrel super-secondary structure. In particular, the sidechains of the pore lining residues maintained their relative orientation during the MD simulation. As illustrative examples, we report a set of representative distances between the amino C-atom of Lys65 and the amide C-atoms of Gln63, between the amino C-atom of Lys65 and the amide C-atoms of Gln61 sidechains and between the amide C-atoms of two neighboring Gln63 sidechains in **Fig. 4**. Steady profiles around ~5 Å and ~6 Å are obtained for the distance between the Lys65 and the Gln63 sidechains and between the Lys65 and the Gln61 sidechains, respectively. Larger fluctuations are observed between the Gln63 residues from 50 ns to 150 ns of the simulated trajectory, but a stationary state around ~4 Å is observed before and after that time window. Conversely, in Pore II this network of interactions is not found due to the location of the Gln61, Gln63, Lys65 residues of interacting monomers at opposing sides of the channel.

**Figure 3.**
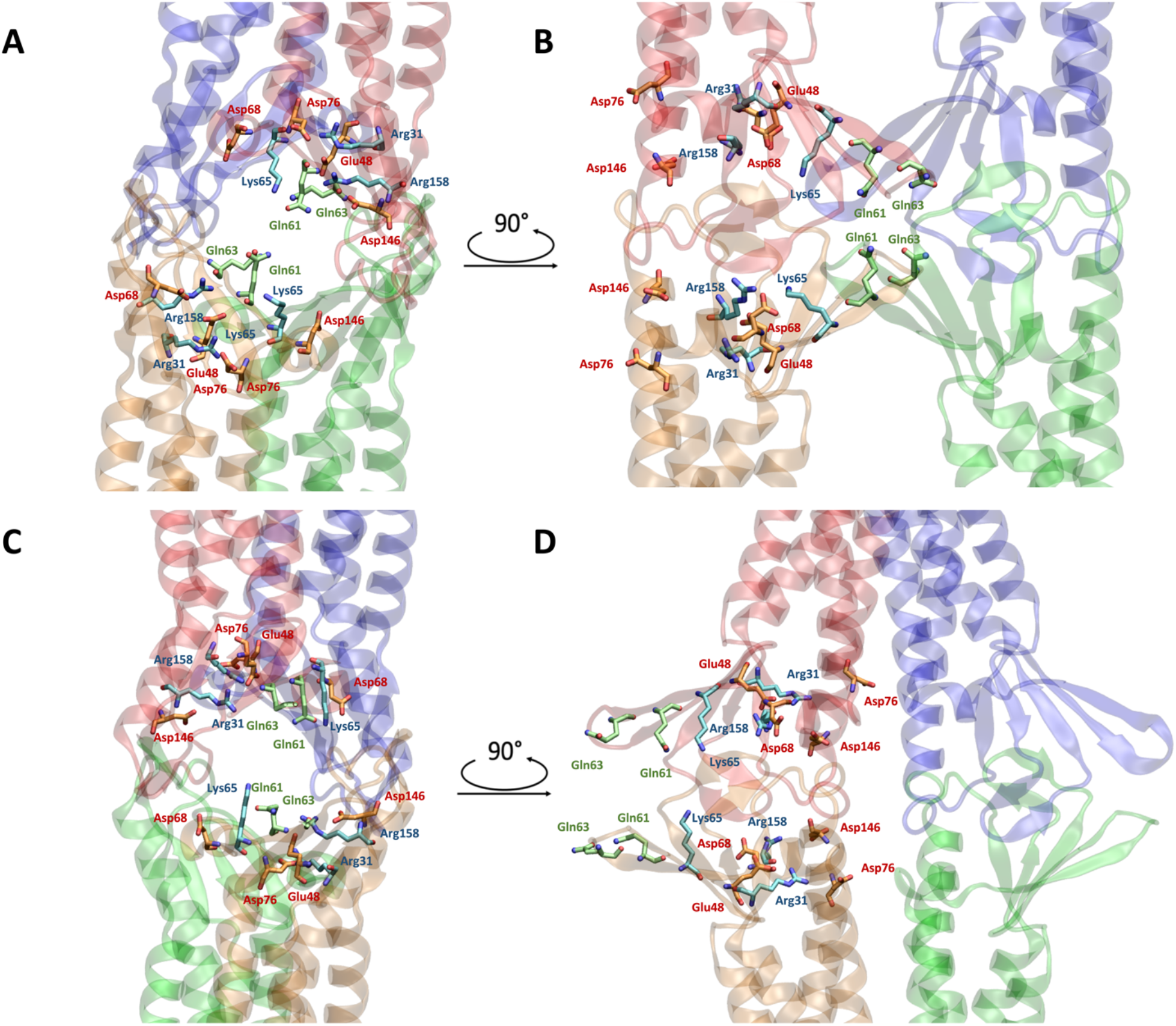
Representations of relevant residues in the two pore models. Apical/basolateral (A) and lateral (B) views of the Pore I configuration. The *β*-barrel arranged by the ECLs of the 4 protomers is visible. Apical/basolateral (C) and lateral (D) views of the Pore II configuration. The reverse orientation of the pore-lining residue compared to the Pore I is visible. The pore-lining residues are indicated for two opposing protomers with respect to the paracellular plane. Acidic residues are depicted in orange, basic residues in light blue and neutral residues in green. Oxygen and nitrogen atoms are shown in red and blue, respectively.

**Figure 4.**
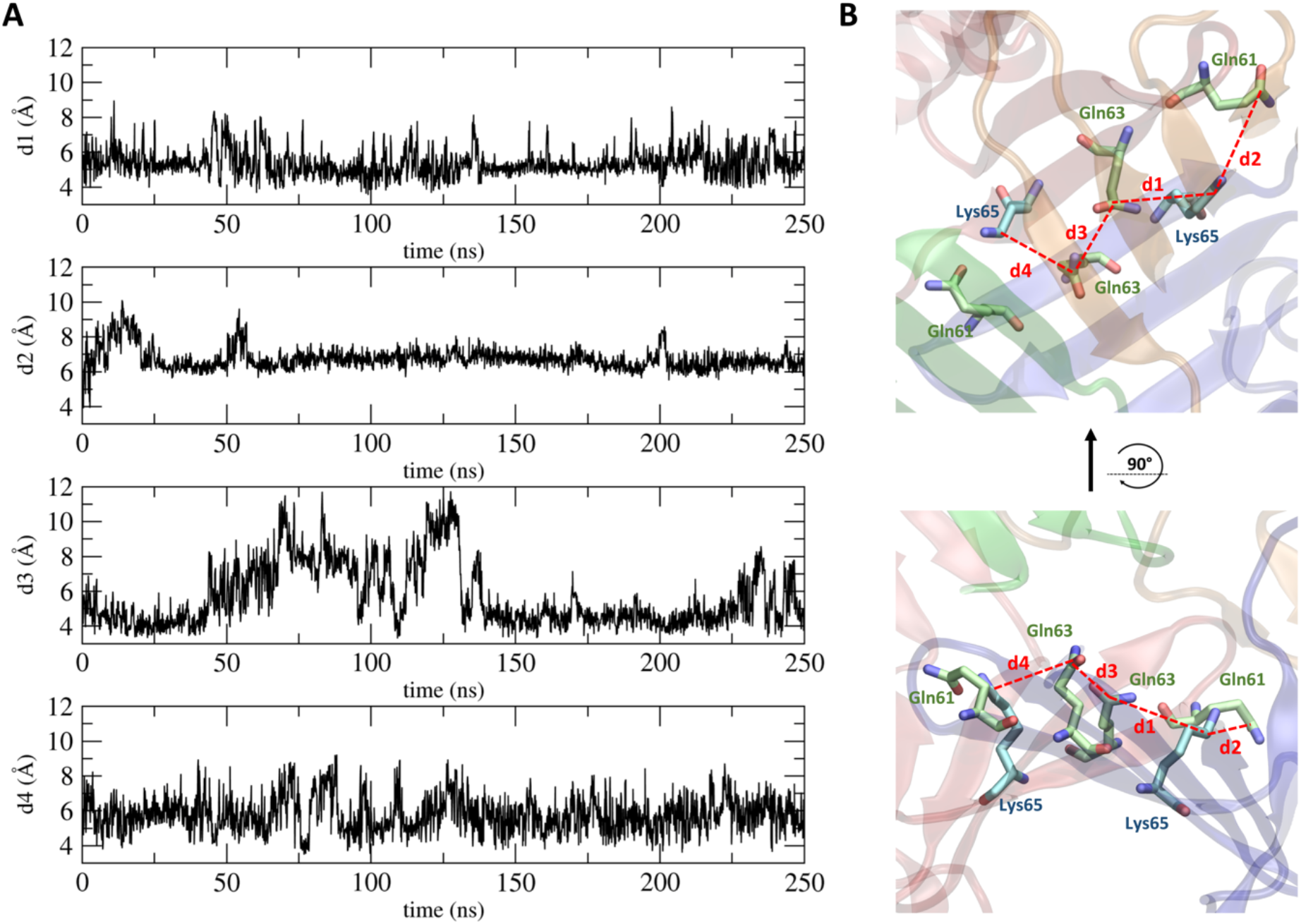
Distances between residues in the central region of the Pore I model. A) Time evolution of distances between the amide C-atom of the Gln residues and the amino C-atom of the Lys residues. B) 3D representation of the computed distances.

### Free energy and electrostatic potential calculations

The physiological roles of Cldn-based TJ pores are differentiated via their capacity to form channels or barriers to the passage of ions and molecules. In this work, we investigated the permeation of water and physiological ions through the two Cldn4 structures. FE profiles were calculated for both the systems using the US method^82^. Results are shown in **Fig. 5** and **Fig. 6**, for Pore I and Pore II, respectively. In these plots, the positions of the pore-lining residues along the pore axis are overlaid to the FE profiles. In the Pore I model (**Fig. 5**), a flat profile is obtained for the passage of the water molecule. On the contrary, barriers of ~4 kcal/mol and 7-8 kcal/mol are observed for monovalent and divalent cations, respectively, all with a symmetrical shape with respect to the pore center, where the Gln61 and Gln63 sidechains are located. In this region, surrounded by the pore-lining Lys65 side chains pointing towards the pore lumen (**Fig. 3A,B**), the pore radius reaches a minimum value (~3 Å), as illustrated in **Fig. 7A**. The four positively charged sidechains are symmetrically paired with respect to the cavity center, with two residues at ~25 Å and two at ~37 Å along the pore axis, generating a positively charged region in the inner section, as revealed by the electrostatic surface reported in **Fig. 8A**. In contrast, the FE profile of the chloride is characterized by a minimum, ~2.5 kcal/mol deep, with inflection points between the Arg31 and Arg158 residues, located close to the pore entrances with respect to the central glutamines (**Fig. 3A,B**).

**Figure 5.**
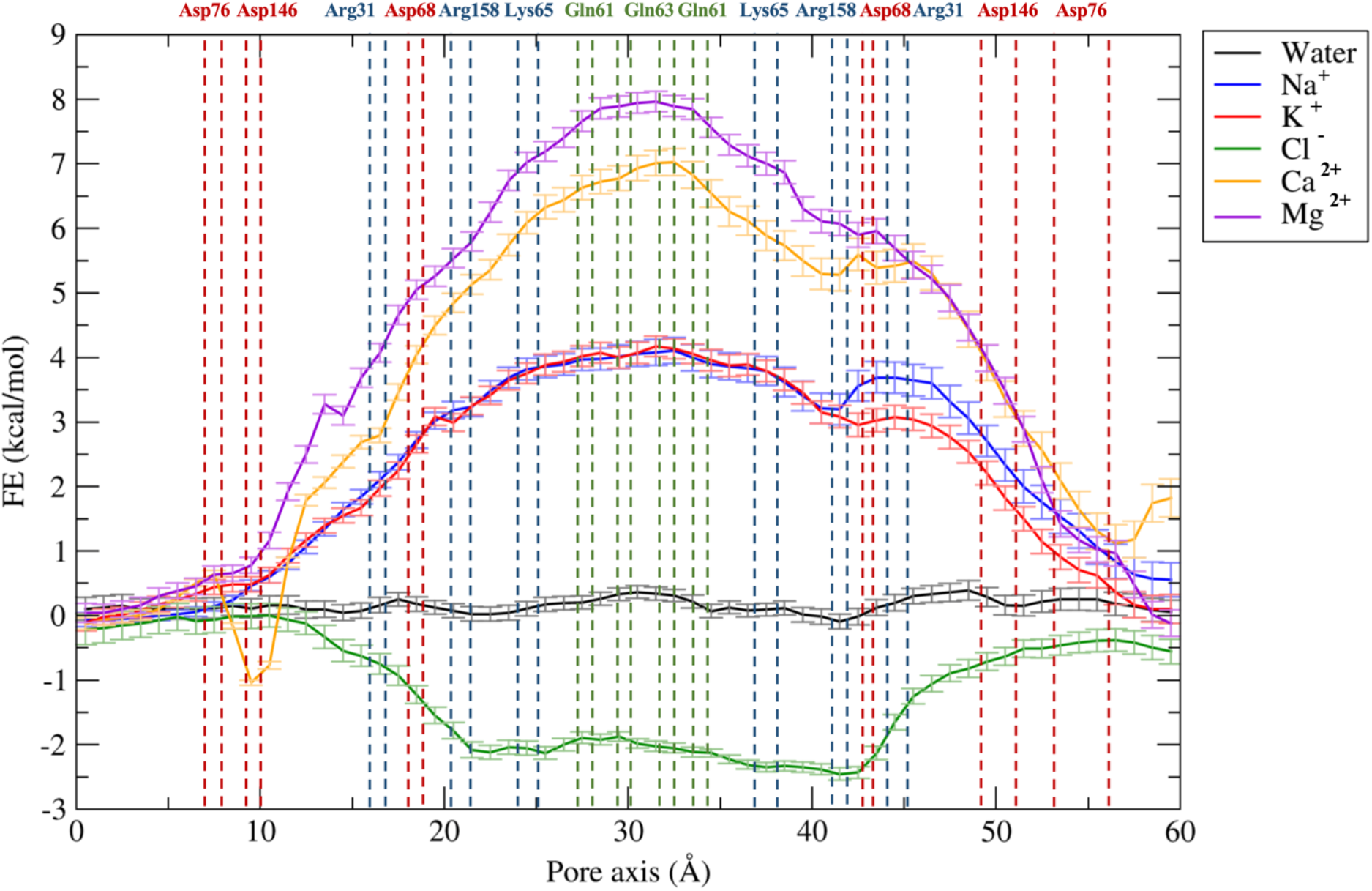
Free energy profiles for the permeation of water and ions through the Pore I model. The position of pore-lining residues along the pore axis coordinates is indicated as dashed vertical lines. Acidic residues are colored in red, basic residues in blue and neutral residues in green.

**Figure 6.**
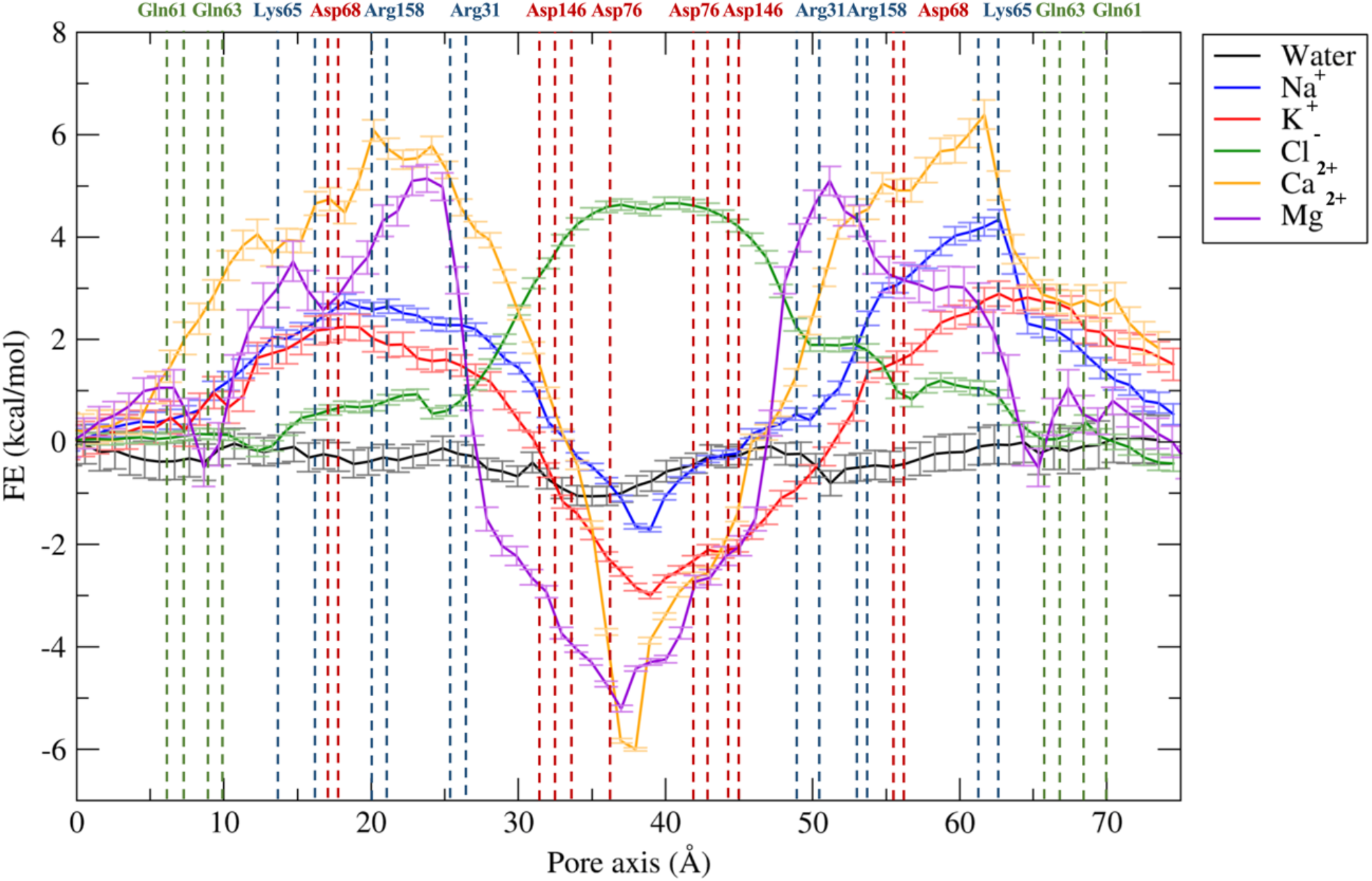
Free energy profiles for the permeation of water and ions through the Pore II model. The position of pore-lining residues along the pore axis coordinate is indicated as dashed vertical lines. Acidic residues are colored in red, basic residues in blue and neutral residues in green.

**Figure 7.**
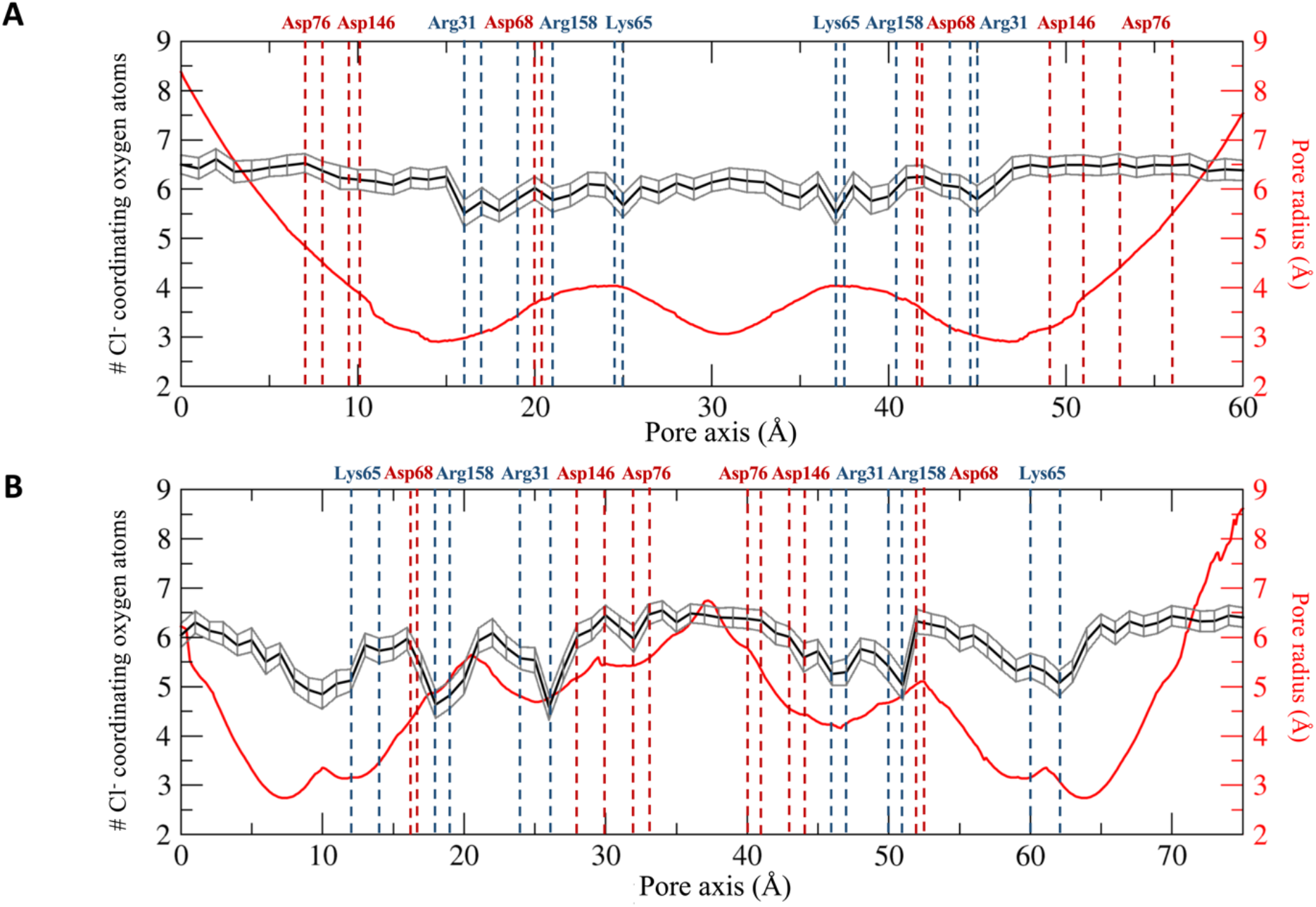
Hydration patterns of chloride and pore radius profile for the Pore I (A) and the Pore II (B) models. The position of the porelining residues driving the ion selectivity is indicated. Acidic residues are shown in red and basic residues in blue.

**Figure 8.**
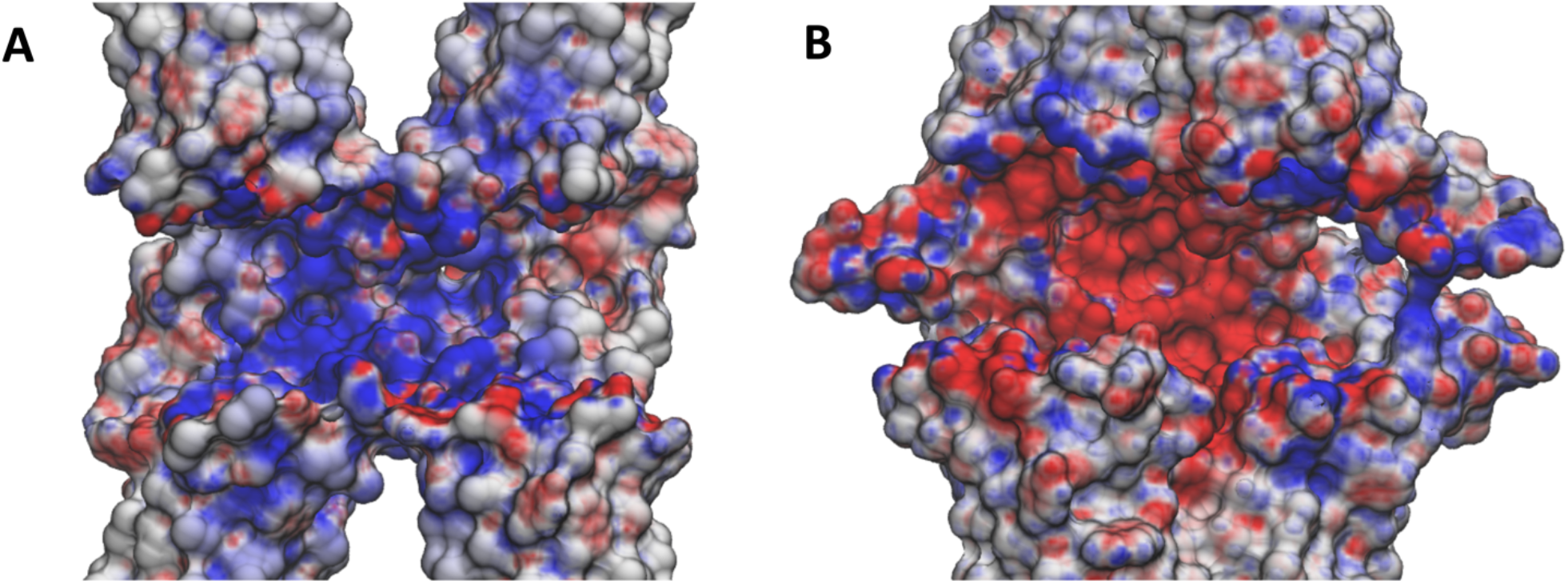
Electrostatic potential surface of the Claudin 4 tetrameric models. Cross sections of the the Pore I (A) and Pore II (B) electrostatic potential surfaces in the paracellular region. The potential surface is shown with a red-white-blue color map with values ranging from −5 to +5 kT/e.

In the Pore II structure, the FE profile for water permeation is also flat (**Fig. 6**). Conversely, all the profiles for cations show two peaks localized at the opposite entrances. Compared to the single peaks obtained for the Pore I model, smaller barriers oppose to the passage of both monovalent (~2.5-3 kcal/mol) and divalent ions (~5.5-6 kcal/mol). The two pairs of Lys65 residues are now close to the pore boundaries (**Fig. 3C,D**), at ~15 Å and ~62 Å along the pore axis, but their position still correlates with the features of the FE profiles. Furthermore, the two FE peaks are located at the narrowest region of the channel, where the radius is at its minimum value (~2.5 Å, **Fig. 7B**). Between the two FE maxima, the profiles for the cations show minima at the cavity center due to the presence of a wide negatively charged chamber formed by the Asp146 and Asp76 residues, consistently with the electrostatic potential shown in **Fig. 8B**. In this region, the Pore II configuration exhibits its maximum pore radius (~6 Å, **Fig. 7B**) and the electrostatic environment is defined by the acidic sidechains pointing towards the lumen of the channel and located between 32 Å and 45 Å along the pore axis (**Fig. 3C,D**). The FE profile of the chloride shows a symmetrical barrier of ~5 kcal/mol at the cavity center, correlating with the positions of the negatively charged side chains of Asp76 and Asp146 (**Fig. 5**). Remarkably, in Pore I the acidic Asp76 and Asp146 residues do not obstacle the chloride flux because they are positioned close to the entrances, and thus have minimal interactions with the anion, as shown by the less extended acidic surface revealed by the electrostatic calculations (**Fig. 8A**). In contrast, the anion passage is mainly driven by the inner, positively charged Arg31, Arg158 and Lys65 sidechains (**Fig. 3A,B**), that are the major determinants of the Pore I cavity electrostatic potential.

### Hydration scheme of the chloride ion

As a further investigation of the mechanisms associated with ion permeation through the two pore models, we computed the average coordination number of chloride in each US window, using a threshold radius of 3.5 Å^88–90^. Results are reported in **Fig. 7** as a function of the pore axis coordinate. The calculated hydration profiles reveal fluctuations in the number of coordinating water molecules surrounding the anion due to pore size variations and contacts with the charged pore-lining residues in the two models. In Pore I (**Fig. 7A**), whose radius varies smoothly and with limited differences in the inner region, minimal variations in the chloride hydration sphere are observed where, on average, the anion loses half water molecule in correspondence of the Arg31, Arg158 and Lys65 positions. The stabilizing interactions with the positively charged pore-lining residues fill the slight depletion of the solvation sphere of the ion and favor its passage through the Pore I cavity, consistently with the energetic minima observed in **Fig. 5**. Conversely, in the Pore II configuration, a more pronounced dehydration of the anion is observed. Chloride loses up to two water molecules in correspondence of the tight regions, where interactions with the sidechains of the positively charged residues (Lys65, Arg31, Arg158) occur. On the contrary, in the inner part of the pore, the anion is fully hydrated, as a consequence of its passage through the widest region of the cavity and the occurrence of unfavorable interactions with the negatively charged pore-lining residues Asp146 and Asp76 (**Fig. 7B**), responsible for the FE barrier, hindering the permeation of the anion in the Pore II model (**Fig. 6**).

## DISCUSSION

The study of the ionic selectivity is a pivotal task to unravel the physiological function of biological channels. While in the literature there is a consolidated state-of-art for the computational investigation of transmembrane ion channel selectivity^91–99^, the study of paracellular channels is still limited to few works ^37,39,40^. In this work, we compared the features of two different paracellular pore models, named as Pore I and Pore II, as putative TJ arrangements formed by Cldn4 proteins. In both the configurations, two copies of Cldn4 dimers, belonging to two neighboring cells, interact with each other in the paracellular space. The Pore I architecture was postulated in the work of Suzuki et al.^41^, describing a homophilic strand of Cldn15-based TJs starting from the crystal structure of the isolated protomer (PDB ID: 4P79)^42^. Notably, it was also proposed independently by Irudayanathan et al.^39^ for Cldns 3 and 5 from docking dimers spontaneously formed in CG MD simulations, first obtained for Cldn5^44^. This configuration was further investigated and refined in various studies for Cldn15^37,38,40,43^, Cldn2^35,36^ and Cldn5^36^. Conversely, the Pore II configuration was first introduced for Cldn5^39^, again by docking dimers assembled in CG MD simulations^44^. Comparisons of Pore I and II conformations were also made for Cldn10b and Cldn3^46^, and for Cldn2 and Cldn4^100^, based only on structural models, with no refinement by MD simulations. In particular, the study in Ref. 100 identified Lys65, Asp68 and Arg158 as key residues for Cldn4 anion selectivity.

Here, we used MD simulations and FE calculations to investigate the reliability of the two putative configurations as pore structures for Cldn4, and to further evaluate their transferability among different Cldn homologs in terms of preservation of tissue-specific physiological functions, governed by the non-conserved ECL1 residues.^30,101^ From a structural point of view, the two models exhibit significant differences at the intracellular, *cis-*interfaces. In the Pore I architecture, two Cldn4 protomers of the same cell interact via a highly hydrophilic interface defined by the Gln61, Gln63 and Lys65 residues in the ECL1 domains (**Fig. 1A**). Oppositely, the dimers originating the Pore II configuration are stabilized by a TM hydrophobic pattern, involving a leucine zipper supported by the *π* - *π* interacA single gene producttions provided by the TM3 Trp138 residues (**Fig. 1B**). As previously described in Ref. 39 for Cldn5, although similar *trans*-interactions are observed in both the models, they lead to opposite arrangements of the pore-lining residues. The Pore I model is characterized by a central region formed by Gln61 and Gln63 pairs, surrounded by the two pairs of Lys65 residues symmetrically positioned with respect to the center, which contribute to the formation of a positively charged electrostatic surface (**Fig. 8A**). Conversely, the Pore II residues arrangement generates an alternating charge environment with Lys65 close to the pore entrances, and two rings of negatively charged sidechains (Asp76 and Asp146) in the central region, resulting in a predominant acidic region in the inner section surrounded by a moderate basic potential at the pore entrances (**Fig. 8B**). Consistently with the values proposed in Refs. 36,39 for the same pore models of other Cldns, the two configurations display a similar minimum pore radius between 2.5 and 3.0 Å (**Fig. 7**). The Pore I duct shows a smooth profile, identifying two bottlenecks symmetrically paired at 15 Å and 45 Å on the pore axis, and a central constriction with a radius of 3 Å (**Fig. 7A**). In contrast, the Pore II channel is tighter at the extremities and shows a central bulge with a radius of 6 Å (**Fig. 7B**). In order to provide a quantitative assessment of the selectivity for the two configurations, we performed US simulations ^82^ to calculate the one-dimensional FE for the permeation of water and physiological ions through the two channels. In the Pore I model, we observed a FE barrier to the passage of cations located at the cavity center with magnitude proportional to the ionic charge (**Fig. 5**). The barriers encountered by the cations are located in the central region where glutamines Gln61 and Gln63 are surrounded by the two pairs of Lys65. The overlap of the FE profiles of cations with same charge indicates a major role of the electrostatic interactions with respect to the steric effects in driving the passage of ions through the channel, similarly to what previously suggested by Alberini et al.^38^ for the Cldn15 cation-selectivity. In contrast, chloride is attracted through the channel by a ~2.5 kcal/mol minimum spanning a region comprised between Arg31 and Arg158, and including the two pairs of Lys65 (**Fig. 5**). Moreover, it was previously suggested that the passage of ions through the narrow region of cation selective Cldn15^38,102^ and Cldn2^82^ pores is associated with the loss of one or more hydrating water molecules. Here, analysis of the hydration state of the anion along the permeation pathway revealed a slight dehydration in correspondence of the positively charged pore-lining residues, able to compensate for the missing water interactions (**Fig. 7A**). On the contrary, the Pore II configuration shows barriers to the passage of all the ions. The pore-lining residues sequence highlights a possible role of the Lys65 residues in reducing the permeation of cations at the pore entrance. The barriers associated to the cations in this pore arrangement are ~1.5-2.0 kcal/mol, lower than those observed in Pore I, but symmetrically located at the two pore mouths. A pronounced minimum is found at the center of the FE profile corresponding to the position of the rings formed by the Asp76 and Asp146 residues that generate a negatively charged chamber with a diameter of almost 12 Å. Consistently, a FE barrier of 5 kcal/mol is obtained at the cavity center for the permeation of chloride (**Fig. 6**). The hydration profile of the anion significantly varies along the cavity. In particular, the particle sheds almost 2 water molecules in correspondence of interactions between the ion and the positively charged sidechains of the Lys65, Arg31 and Arg158 residues and where the pore reveals major constrictions. Conversely, it preserves its hydration sphere in the central region of the cavity where the unfavorable contact with the negatively charged residues takes place (**Fig. 7B**) and the maximum in the FE profile is observed (**Fig. 6**). This evidence confirms the driving role of electrostatic interactions in the formation of the FE barriers for the anionic permeation.

In conclusion, the two pore models suggest opposite behaviors for the Cldn4-based TJ functionality, hence resulting mutually exclusive. Remarkably, the Pore I configuration provides the first atom-detailed model recapitulating the observed anion-selectivity of Cldn4 based TJs ^23–25,27^. In contrast, the Pore II model forms a barrier for both cations and anions^28,32,101,103,104^. The physiological role of Cldn4 as channel or barrier remains elusive^23–28,30,32,33,101,103–105^, and the factors modulating the expression of its functionality have not been fully clarified yet^30^. The capability to arrange in strands is also questioned due to the dependence on the cell expression system^26,30–32,105^, although experimental data supporting it have been reported^106^. Notwithstanding, our FE profiles for the Pore I configuration show higher barriers to the passage of the cations than Pore II. According to these considerations, we propose Pore I as the putative configuration to recapitulate the physiological properties of anion-selective Cldn4-based TJs. Further experimental characterization of Cldn4 physiology will ultimately provide the necessary knowledge to discriminate between different predicted configurations of Cldn4-based TJs. Additional structural studies are also required to refine the peculiarities of the two architectures discussed in this work. In particular, the atom-detailed characterization of Pore II model is currently less complete than Pore I model, and fewer experimental results supporting this arrangement are available to date^45^. Testing these models for different Cldns in terms of tissue-specific TJ physiology will provide a fundamental contribution to their validation.

## Author contribution

A. Berselli contributed to design the research, performed the simulations, analyzed the data, participated to the discussion for the data interpretation and wrote the manuscript. G. Alberini shared his know-how for the simulations set-up, supervised the data production and analysis, participated to the discussion and wrote the manuscript. L. Maragliano and F. Benfenati supervised the project, discussed the results and revised the manuscript.

## Acknowledgments

We are grateful for the HPC infrastructure and the Support Team provided by the Fondazione Istituto Italiano di Tecnologia (IIT) that allowed us to run our MD simulations. We acknowledge Andrea L. Benfenati, Sergio Decherchi and Diego Moruzzo for valuable help. We thank Jörg Piontek and Matteo Ceccarelli for useful discussions. The research was supported by IRCCS Ospedale Policlinico San Martino (Ricerca Corrente and 5×1000 grants to FB and LM) and by Telethon/Glut-1 Onlus Foundations (seed project 1754 to FB).

